# Imaging transmembrane dynamics of biomolecules at live cell plasma membranes using quenchers in extracellular environment

**DOI:** 10.1101/2020.08.30.266148

**Authors:** Wenqing Hou, Dongfei Ma, Xiaolong He, Weijing Han, Jianbing Ma, Chunhua Xu, Ruipei Xie, Qi-hui Fan, Fangfu Ye, Shuxin Hu, Ying Lu, Ming Li

## Abstract

It is a big challenge to measure position changes of biomolecules in the direction normal to the plasma membranes of living cells. We developed a one donor-multiple quenchers Fӧrster resonance energy transfer method by using non-fluorescent quenchers in the extracellular environment. It senses subnanometer position changes of a fluorophore-labeled biomolecule in the plasma membrane. The method was validated by monitoring flip-flops of individual lipid molecules incorporated in plasma membranes. We studies membrane perforation by a host defense peptide from the extracellular side and found that the pore-forming peptide is dynamic, switching among different insertion depths. The method is especially useful in studying interactions of membrane proteins with the inner surfaces of plasma membranes. Our method will find wide applications in systematic analysis of fundamental cellular processes at plasma membranes.

## Introduction

Plasma membranes are permeable barriers between cells and their environment. They control the flow of information and the trafficking of substances in and out of cells, and have become the target of various drugs in the process of curing diseases (1). Many important physiological processes, such as receptor-ligand interactions, lipid flip-flops and lipid-raft formation, are accompanied with the motions and structural changes of biomolecules on plasma membranes (2, 3). Current techniques such as antibody labeling and tracking and in-cell NMR have produced vast valuable findings in this field (4-6). However, advancing the research of membrane-associated processes still demand technologies able to measure in real time positional and conformational changes of biomolecules at live cell plasma membranes (7). It is a substantial challenge because cellular membranes are sophisticated matrices composed of thousands of biomolecules (8). This is especially true when positional changes in the direction normal to the membranes are the main concern (6). One needs to extract useful information from complicated movements of the biomolecules and the membranes. The biomolecules undergo in-plane diffusions in the membranes, which in turn undergo strong out-of-plane fluctuations with amplitudes usually exceeding their thicknesses. Fluorescence imaging and spectroscopic techniques are often the methods of choice for investigating membrane proteins in live cells. Conventional fluorescent resonance energy transfer (FRET) probes angstrom-scale movements of donor-labeled proteins (9, 10). It is however only useful when the positions of donors or acceptors are precisely known. These are not easily determined because both the donors and acceptors diffuse in fluidic membranes (11, 12). In addition, when fluorophore-labeled molecules form clusters with unknown sizes and conformations, it is often difficult to correctly convert FRET efficiency to inter-fluorophore distance (13).

In our previous work, we developed an in vitro fluorescence method to measure positions of membrane proteins in lipid bilayers (14). The method enabled us to extract information about the transmembrane motions of biomolecules while disregarding their in-plane motions in simple liposomes. However, complex feedback networks and/or molecular crowding in cells may alter the functions and dynamics of biomolecules, potentially posing obstacles to the physiological relevance of information gained from the in vitro systems. The in vitro approach therefore cannot be directly applied to study the trafficking of biomolecules in plasma membranes of live cells because it is very hard to deliver high concentration exogenic substances into live cells. In the present work, we extend our approach to live cells by adding quenchers in the extracellular environment during cell imaging. The feasibility of the approach relies on appropriate selection of the quenchers. They should be neutral, water soluble and biocompatible. Most importantly, they should not adsorb onto the cell surfaces specifically or non-specifically. The strategy enabled us to monitor positional changes of fluorophore-labeled biomolecules relative to the plasma membrane surfaces. We anticipate that the method will significantly improve researches on dynamics of membrane proteins, responses of receptors, signal transductions across plasma membranes and so on, which are not assessable by conventional technologies.

## Results

### Design and principle of the method

The underlying mechanism of our method is FRET between a single fluorophore and a multitude of quenchers in the extracellular environment (termed queenFRET hereafter) (Fig. 1*A*). The method does not require hardware modification to conventional fluorescence imaging microscopes. It works well with total internal reflection fluorescence microscopy (TIRF) and fluorescence lifetime imaging microscopy (FLIM). The FRET between the fluorophore and the quenchers results in a very steep change in fluorescence as the fluorophore-labeled molecule moves from the outer to the inner surface of the membrane or vice versa. The calculated FRET efficiency *E* as a function of distance between the fluorophore and the quencher solution is presented in the form of relative fluorescent intensity or relative lifetime in Fig. 1*B*, that is, *F/F*_*0*_ *= τ/τ*_*0*_ *= 1-E*. As a result, the positional changes of the fluorophore-labeled biomolecule in cell membranes can be reflected by its intensity or lifetime changes (Fig. 1*C*).

**Fig. 1.**
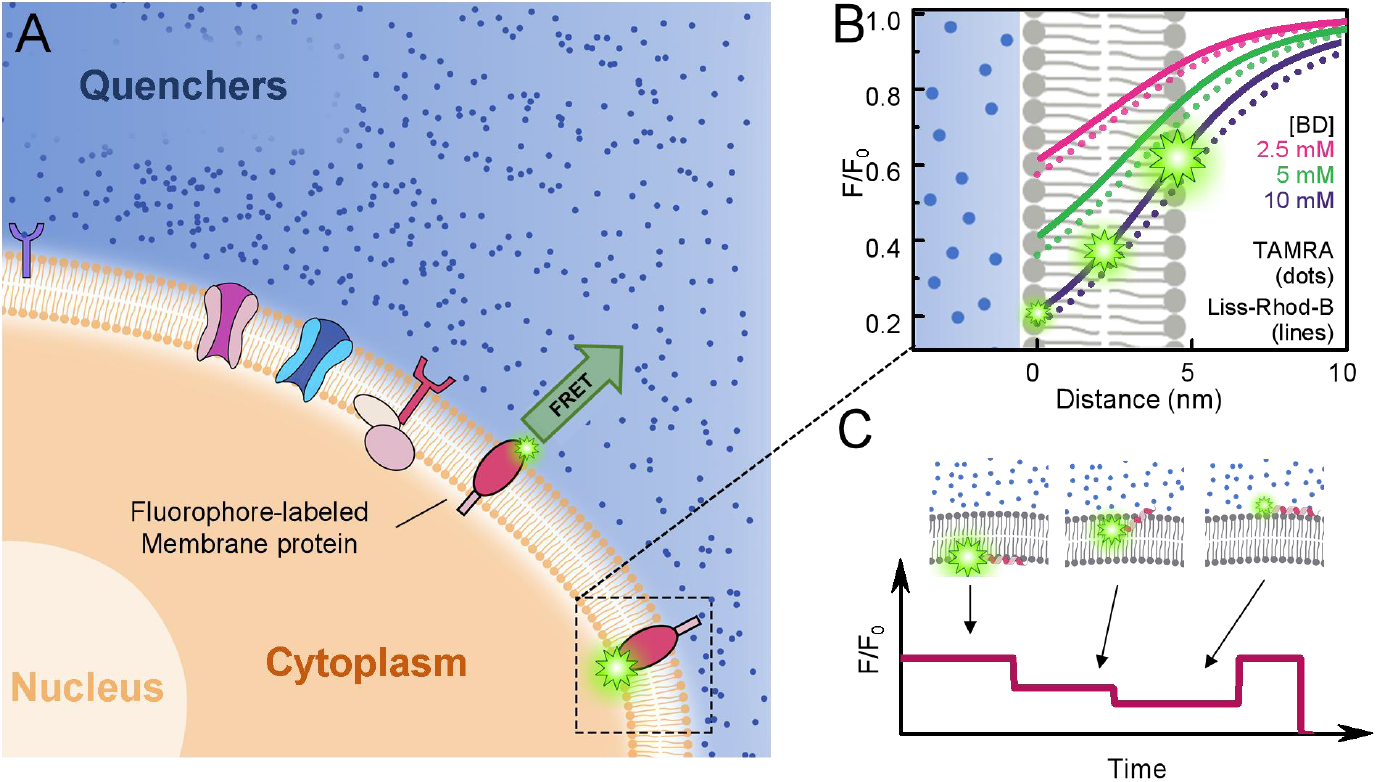
Principle of queenFRET. (*A*) Scheme of the method, in which the fluorescent energy is transferred from single fluorophores at the membrane to multiple quenchers in the extracellular environment. (*B*) Calculated *F/F*_*0*_ or *τ/τ*_*0*_ vs distance at various concentrations of the quenchers. (*C*) Scheme of a time course of the fluorescent intensity when the fluorophore transits from the inner surface to the outer surface of a cell membrane.

In practice, one should always measure the intrinsic fluorescent intensity *F*_*0*_ and the intrinsic fluorescence lifetime *τ*_*0*_ of the donors in parallel with *F* and *τ*. On one hand, the measurements in the absence of the quenchers can function as a control to ensure that any changes in the fluorescent intensity are induced by the changes in the distance. On the other hand, many fluorophores may display different photo-physical properties in different environments. Such environment-sensitive fluorophores should be excluded from the candidates of the donors if they displayed fluorescent fluctuations in the absence of the quenchers. Those that are fluorescently unstable, e.g., blinking frequently, should also be excluded. The quencher used in the present study was blue dextran (BD), although we believe that many other dyes with appropriate absorption spectra may be also effective. BD is a high molecular weight dextran with attached Cibacron Blue whose absorption maximum is ∼620 nm (15). BD is used in various fields such as pharmaceutical, photographic and agricultural industries owning to its favorable properties, namely neutrality, water solubility, and biocompatibility (16). In our experiments, the cells remained viable even after incubation in BD-containing culture medium overnight.

### Flip-flops of lipid molecules in plasma membranes

We first tested the feasibility of our method by measuring flip-flops of fluorophore-labeled lipids. Flippases often translocate aminophospholipids, e.g., phosphatidylserine (PS) and phosphatidylethanolamine (PE), towards the cytosolic leaflet of plasma membranes to maintain their asymmetry (17-22). We adopted a vesicle-plasma membrane fusion method (23-25) to incorporate fluorescent PE (Liss-Rhod-B-PE) into the membrane. When a cell adheres to a glass surface, there is usually a gap between the basal membrane and the surface, which typically varies between 20 and 100 nm depending on the cell type (26). Therefore, a pseudo total internal reflection (pseudo-TIR) mode (27, 28) in which the incident angle is set slightly smaller than the critical angle for TIR was adopted to avoid intensity variation induced by the evanescent field along the z-axis (Fig. 2*A*). Individual fluorophores were tracked and the intensities of the fluorophores were recorded (Fig. 2*B*). In the absence of BD, the fluorescence of the fluorophores was stable (Fig. 2*C*). The BD solution (5 mM) percolated into the gap to quench the fluorophores in the membrane. Because the fluorophores in the two opposite surfaces of the lipid bilayer have different distances to the quencher solution, they displayed two relative fluorescent intensities (Fig. 2 *E* and *F*) with *F/F*_*0*_ = 0.38 ± 0.07 (mean ± s.d.) for the outer surface (cyan) and 0.80 ± 0.07 for the inner surface (dark cyan), where *F*_*0*_ was the intrinsic fluorescence in the absence of the quenchers (Fig. 2*D*). The centers of the two peaks represent a difference in distance by 5.2 ± 0.9 nm in consistence with the thickness of the plasma membrane (8, 29, 30), indicating that queenFRET is precise enough to measure the location of a single fluorophore in plasma membranes. Interestingly, beside the traces that displayed low or high intensities, we also observed many traces that illustrated flipping of the fluorophore-labeled PE from the outer leaflet (with low intensity) to the inner leaflet (with high intensity). To the best of our knowledge, this is the first observation of single lipid molecules flipping in live cell membranes.

**Fig. 2.**
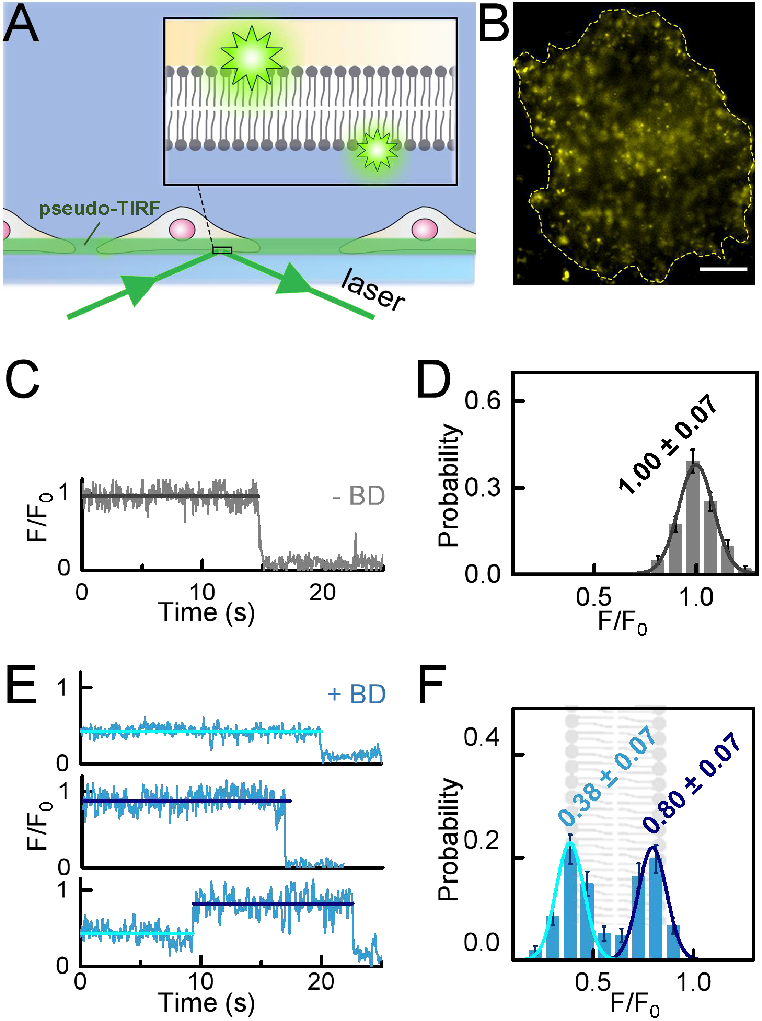
TIRF-based analysis of lipid molecules in the inner/outer leaflets of the cell membrane. (*A*) Scheme of the pseudo-TIRF illumination on adherent cells. (*B*) Image of the basal membrane containing Liss-Rhod-B-PE. Scale bar in the image is 10 μm. (*C*) Typical fluorescence trace and (*D*) the corresponding intensity histogram of Liss-Rhod-B-PE in the absence of the quenchers. (*E*) Typical fluorescence trace and (*F*) the corresponding intensity histogram of Liss-Rhod-B-PE in the presence of 5 mM BD in the culture medium. Cyan and dark cyan traces and histograms represent lipid molecules at the extracellular and the intracellular surfaces, respectively.

The positions of the fluorophores in the membrane can also be deduced from their fluorescence lifetimes measured with FLIM by using a time-correlated single photon counting (TCSPC) system (Fig. 3 *A* and *B*). Single molecules cannot be readily imaged in this mode. However, the lifetimes of the fluorophores in a pixel (with a 1-ms dwelling time) can be deduced by fitting the exponentially decaying curve (the TCSPC curve), provided that the fluorophores are sparse enough so that there are less than two fluorophores in each pixel on average (see details in SI). The lifetime images of the plasma membranes looked differently in the absence and presence of BD (Fig. 3*B*). We divided the images into small units and calculated the lifetime of each unit to obtain lifetime histograms). The histogram of the intrinsic lifetimes of Liss-Rhod-B attached to PE displayed a peak at 2.8 ± 0.2 ns (Fig. 3*C*, upper panel). When BD solution (5 mM) was added to the petri dish mounted on the microscope, two additional lifetimes were observed: 1.0 ± 0.2 ns for the outer surface and 2.3 ± 0.2 ns for the inner surface (Fig. 3*C*, lower panel). We then adopted time-lapse imaging to follow the transmembrane transport of the lipids in Fig. 3*D*. A large proportion of the Liss-Rhod-B-PE molecules were transferred to the inner leaflet when the fluorescent lipids had been incorporated into the membrane for about 120 min. In contrast, no change was observed for the phosphatidylcholine (PC) lipids (Fig. S4 in SI Appendix).

**Fig. 3.**
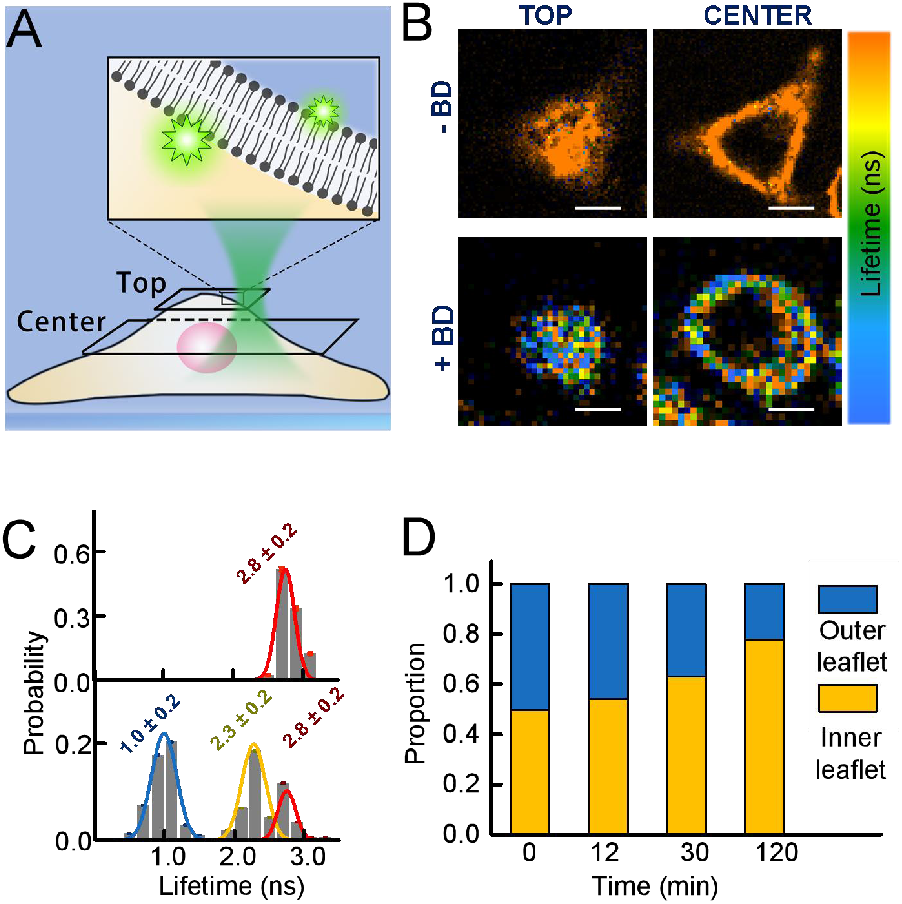
Confocal-based analysis of lipid molecules in the cell membrane. (*A*) Scheme of the confocal-based FLIM imaging. (*B*) FLIM images of the top-(left) and the mid-plane (right) of the cell in A. The two upper panels are images in the absence of the quenchers. The two lower panels are images in the presence of 5 mM BD in the culture medium. (*C*) Lifetime histograms of the fluorophores in B in the absence (upper panel) and presence (lower panel) of the quenchers. (*D*) Proportions of the Liss-Rhod-B-PE molecules in the outer leaflet (blue) and inner leaflet (yellow) according to the time-lapse imaging of the cells.

### Interaction of extracellular proteins with plasma membranes

We next demonstrated that queenFRET can monitor interactions of extracellular proteins with plasma membranes. We labeled the human host defense peptide LL-37 with tetramethylrhodamine (TAMRA) at its N-terminus (TAMRA-LL-37) and added the peptide to the culture medium to monitor its membrane insertion dynamics in the lung epithelial carcinoma cell line A549 (Fig. 4). LL-37 is a positively charged cathelicidin that targets negatively charged bacterial membranes (31). It also binds cancer cells with exposed anionic lipids (32). Upon incubation, the peptide percolated into the thin gap between the basal membrane and the glass surface, where it interacted with the plasma membranes (Fig. 4*A*). After illuminating the sample for 30 s to bleach a large fraction of the TAMRA-LL-37 molecules, individual fluorescence spots were observed to diffuse on the basal membrane. At a low protein concentration (1 nM), LL-37 usually maintains as monomers, and typical intensity traces showed a single photo-bleaching step (Fig. 4*A*, left panel; Fig. 4*B*, grey trace). At higher protein concentrations, for example at 20 nM LL-37, some bright fluorescence spots could be observed (Fig. 4*A*, right panel). Typical intensity traces of these spots displayed multiple photo-bleaching steps, indicating that LL-37 formed oligomers at higher concentrations (Fig. 4*B*, cyan trace). Fig. 4*C* illustrates the comparison of the monomeric TAMRA-LL-37 (1 nM) in the absence (grey) and presence (pink) of 5 mM BD. Both intensity traces are stable. The ratio of the fluorescent intensities, *F/F*_*0*_, is in agreement with the calculation assuming that the monomeric TAMRA-LL-37 adsorbed and laid on the surface of the A549 cell (Fig. 4*D*). At an elevated concentration (1 nM TAMRA-LL-37 mixed with 100 nM wild type LL-37), LL-37 became dynamic, switching between different depths (Fig. 4*E*). While the intensity of TAMRA-LL-37 remained stable in the absence of BD (grey), the intensity histogram of the dynamic TAMRA-LL-37 displayed three major peaks in the presence of BD (pink). The peaks can be converted to insertion depths of 0.1 ± 0.9, 2.1 ± 0.9 and 4.7 ± 0.9 nm, respectively (Fig. 4*E*). The fourth peak with the highest intensity could be attributed to fluorophores near the intracellular surface, which could not be accurately positioned because they were in the insensitive region of our approach. The results were similar to those obtained from our previously developed in vitro methods (33), again demonstrating the validity of queenFRET. The results suggest that LL-37 forms oligomers and displays dynamic behavior in live cell membranes, possibly associated to the mechanism of its anti-cancer functions (32).

**Fig. 4.**
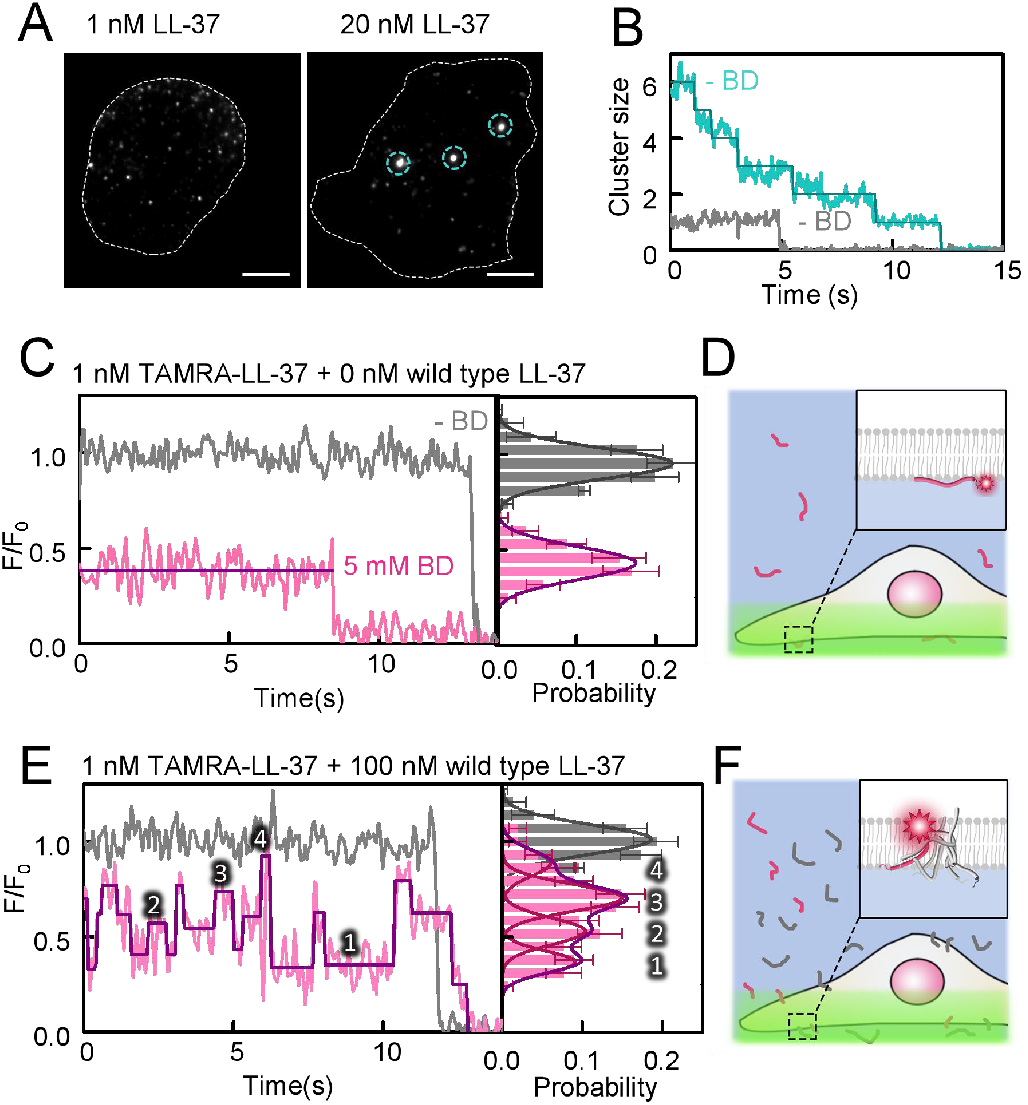
Interaction of the extracellular LL-37 proteins with plasma membranes. (*A*) Pseudo-TIRF images of N-TAMRA-LL-37 at the plasma membranes at low (left) and high (right) concentrations. LL-37 forms clusters at high concentrations as indicated by the cyan cycles. (*B*) Photo-bleaching of an LL-37 monomer (grey) and an LL-37 cluster (cyan) in A. (*C*) Typical single molecule fluorescence traces of LL-37 at low concentrations and the corresponding intensity histogram. (*D*) Scheme of LL-37 at the inner surface of the plasma membrane. (*E*) Typical single molecule fluorescence traces of LL-37 at high concentrations and the corresponding intensity histogram. (*F*) Schemes of LL-37 inside the plasma membrane. Grey data were acquired in the absence of the quenchers. Statistics were from 110 traces of 35 cells.

### Interaction of intracellular proteins with plasma membranes

We finally investigated insertion of proteins from the intracellular side of cell membranes, which is valuable when endogenous proteins are of interest. Conventional techniques were only able to tell whether a protein of interest is at the membrane or not. QueenFRET can not only distinguish different membrane-inserting proteins but also different insertion states of a same protein in the membrane. For simplicity, we delivered purified proteins into the cells to mimic endogenous proteins. The mixed lineage kinase domain-like protein (MLKL) was used as an example, which is crucial for necroptosis by permeabilizing membranes through its N-terminal region upon phosphorylation (34, 35). It was reported that the short form of MLKL (residues 2-123; MLKL_2-123_) is more active than the long form of MLKL (residues 2-154; MLKL_2-154_) which contains a brace helix (19). We labeled MLKL_2-123_ and MLKL_2-154_ at the residue S55 with TAMRA and delivered them into the human monocytic-like cell line THP-1. The cells were then suspended in a microfluidic channel and imaged by FLIM (Fig. 5*A*). The lifetime histogram for MLKL_2-154_ indicated that they inserted into the membrane shallowly (peak at 2.4 ns in Fig. 5*B*). In contrast, MLKL_2-123_ might insert into the membrane more deeply at 2.7 nm (1.9 ns peak) and 4.6 nm (1.4 ns peak), respectively (Fig. 5*C*). The extra peak (red fitting curve) represent molecules in the cytoplasm. Our results are qualitatively consistent with previous reports (36), but queenFRET provides a much more quantitative description. The observation that MLKL_2-123_ inserts into the membrane in three different depths is very interesting. It suggests that the protein may change conformations upon interaction with the membrane, a phenomenon that deserves deeper investigation in the future.

**Fig. 5.**
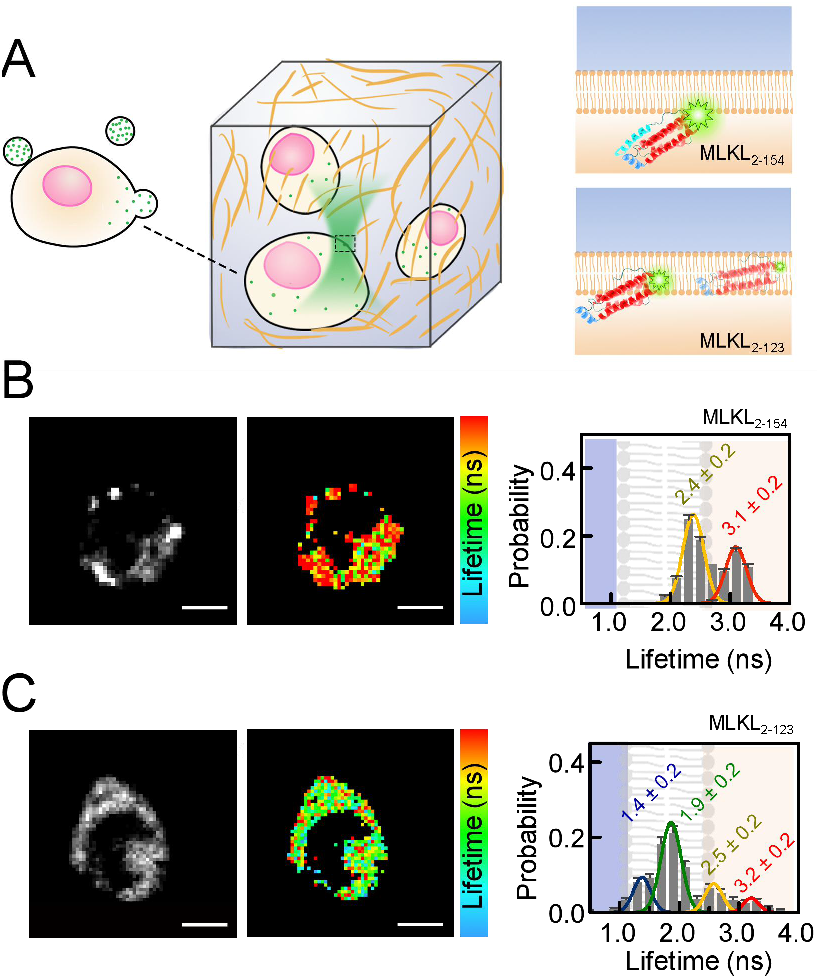
Interaction of the intracellular MLKL proteins with plasma membranes. (*A*) Scheme of MLKL in the THP-1 cells suspended in a microchip for confocal-based FLIM. (*B*) Intensity image (left), lifetime map (middle) and lifetime histogram (right) of TARMA-labeled MLKL_2-154_. (*C*) Data for TARMA-labeled MLKL_2-123_. Color bar is from 1.5 to 3.0 ns. Scale bar = 5 μm. The concentration of BD was 5 mM. The histograms were built from data of 10 cells.

## Discussion

Probing structural dynamics of biomolecules in plasma membranes had been a very difficult task. Here, we demonstrated the feasibility of queenFRET in quantitating transmembrane trafficking of fluorophore-labeled biomolecule from not only the inside but also the outside of live cells. Both locations are crucial to understanding protein-membrane and protein-ligand interactions. Our method is especially useful when the biomolecules of interest interact with the cytosolic leaflet of the plasma membrane, which is not accessible by conventional techniques at the single molecular level. QueenFRET can be implemented with both TIRFM and FLIM. The prerequisite is to label the molecules-of-interest with small fluorophores. TIRFM is preferred when investigating dynamics of plasma membranes adhered to glass surfaces. Confocal-based FLIM is a good choice to observe floating cells such as those in microfluidic channels or even inside a cluster of cells.

Choosing appropriate quenchers and donors is crucial to perform the queenFRET successfully. The quenchers should fulfil the following requirements: **1)** Sufficient solubility. According to our calculations (Fig. 1*B*), the concentration of the extracellular quenchers should be as high as a few millimoles per liter to produce sufficient FRET with the donors; **2)** Appropriate spectral properties. The quenchers, which serve as the FRET acceptors, should have high extinction coefficient. In addition, the absorption spectrum of the quencher should show significant overlap with the emission spectrum of the donors, allowing effective energy transfer even at relatively large donor-quencher distances so that the positional changes of the donor near the cytosolic surface of the cell membrane can be readily observed; **3)** Low fluorescence emission. The quenchers should be non-fluorescent (at least of low emission at the detection window of the donors) in order not to interfere with the fluorescent signals of the donors. Moreover, as in other *in vivo* measurements, additional requests on the quenchers should be satisfied: **1)** Non-toxicity. Biocompatibility is the primary factor to be considered. For example, some dyes may undergo photochemical reactions and cause damages to various biological substrates (37, 38). BD in our work is non-toxic to cells, allowing us to observe biomolecule dynamics for a few hours; **2)** Negligible interactions with cell surface. Plasma membranes are complicated systems that consist of various biomolecules with different physical and chemical properties. Interactions of the plasma membranes with the quenchers may alter their spatial distribution near the membranes, thus change the dependence of FRET efficiency on the distances, resulting in uncertainty when interpreting the fluorescent signals. Ideal quenchers should not have any interactions with the cell surfaces. Care should also be taken when choosing the donors. Fluorophores used to label the biomolecules should avoid strong interactions with the membranes because they may alter the membrane-interacting patterns of the biomolecules of interest (39). In addition, fluorophores that are sensitive to solvent polarity should also be avoided unless the fluorophores remain in the same environment, for example, always in the membrane during the measurements (40).

It is noteworthy that the absolute fluorescence intensity is not necessary to gain the positions of the fluorophores in the membranes. In our approach, the distance (*d*) between the donor and the interface of the quencher solution and the membrane was determined by comparing the measured relative intensity (*F/F*_*0*_) or relative lifetime (*τ/τ*_*0*_) with the calculated curves like the ones displayed in Fig. 1*B*. One should first measure the intrinsic parameters *F*_0_ and *τ*_*0*_ for the fluorophore of interest under the same experimental conditions but the quenchers and use them as the reference. QueenFRET is of sub-nanometer precision. We can estimate the error in distance from the inverse of the digital differentials of the theoretical intensity-distance curves. The same analysis applies to errors in the lifetime measurements.

In the present work, we have focused on the transmembrane positions of fluorophores disregarding their in-plane movements. Our method can work along with many localization-based imaging techniques. For example, the in-plane movements of the biomolecules can be recorded by existing methods such as single-molecule tracking and super-resolution imaging. We anticipate that the combination of queenFRET with super-resolution imaging may significantly improve our understanding of membrane trafficking, receptor responses and signal transductions of live cells, to name just a few.

## Acknowledgments

This work was supported by National Natural Science Foundation of China (Grant Nos. 11974411 to YL, and 91753104 to ML), CAS Key Research Program of Frontier Sciences (Grant Nos. QYZDJ-SSW-SYS014 to M.L. and ZDBS-LY-SLH015 to Y.L.), and CAS Youth Innovation Promotion Association (Grant No. 2017015 to YL). The authors also gratefully acknowledge the support of the K. C. Wong Education Foundation.

